# Myc supercompetitor cells exploit the NMDA receptor to subdue their wild-type neighbours via cell competition

**DOI:** 10.1101/2020.02.11.943498

**Authors:** Agnes R. Banreti, Pascal Meier

## Abstract

Myc is a major driver of cell growth in many cancers, but direct inhibition of Myc’s oncogenic activity has been challenging. Interactions between wild-type and Myc-expressing cells cause Myc cells to acquire ‘supercompetitor’ behaviour that increases their fitness and enables them to overtake the tissue by killing their wild-type neighbours through TNF-induced cell death during a process called cell competition. Here we report that the competitive behaviour of Myc, RasV12 cells, and normal epithelial cells, critically depends on the NMDA receptor. Myc cells upregulate NMDAR2 (NR2) to gain supercompetitor status and subdue their wild-type neighbours. Pharmacological inhibition or genetic depletion of NR2 changes the supercompetitor status of oncogenic Myc or RasV12 clones into ‘superlosers’, resulting in their elimination via cell competition by wild-type neighbours in a TNF-dependent manner. Our data demonstrate that that the NMDA receptor (NMDAR) determines cellular fitness during cell competition, and can be targeted to change the fitness landscape of supercompetitive Myc and RasV12 clones, converting them into superlosers.

## Introduction

Cell competition is an evolutionary conserved quality control process that eliminates suboptimal, but otherwise viable cells, thereby safeguarding tissue fitness and tissue homeostasis^1^. How relative fitness disparities are measured across groups of cells, and how the decision is taken whether a particular cell will persist in the tissue (‘winner cell’) or is killed (‘loser cell’) is not completely understood^2^. This is an important issue as competitive behaviour is exploited by cells with deregulated oncogenes or tumour suppressors, which subsequently expand at the expense of, and with the help from, their cooperating wild-type (WT) neighbours^1^.

Growth signalling pathways involved in metabolic cell competition all seem to funnel through Myc, which functions as an essential signalling hub sustaining growth of many types of cancers. Myc regulates expression of components that control proliferation, cell death, differentiation, and central metabolic pathways. Particularly, acute changes in cellular metabolism appear to be critical for the winner phenotype during Myc supercompetition in *Drosophila*^3^.

Recent data demonstrate that living tumours can uptake lactate and preferentially utilize it over glucose to fuel TCA cycle and sustain tumour metabolism^4^. Moreover, the growth promoting effect of stromal cells is impaired by glycolytic inhibition, suggesting that the stroma provides nutritional support to malignant cells by transferring lactate from cancer-associated fibroblasts (CAFs) to cancer cells^5, 6^. Such energy transfer from glycolytic stromal cells to epithelial cancer cells closely resembles physiological processes of metabolic cooperativity, such as in ‘neuron-astrocyte metabolic coupling’ in the brain, and the ‘lactate shuttle’ in the skeletal muscle^7, 8^. Activation of glycolysis in astrocytes and MCT-mediated transfer of lactate to neurons supports neuron mitochondrial oxidative phosphorylation and energy demand^9^. These observations raise the intriguing possibility that lactate serves as fuel to complement glucose metabolism in both supercompetitors as well as tumours.

We report here that the NMDA receptor, a key component of the ‘learning and memory’ module, is repurposed to coordinate the competitiveness of epithelial cells within a tissue. We find that the activity of the NMDAR underpins cellular fitness in *Drosophila* epithelia during Myc- and RasV12-induced supercompetition, paradigms of cell competition with parallels to mammalian tumour promotion. Inactivation of NMDAR changes the supercompetitor status of oncogenic Myc or RasV12 clones into *‘superlosers’*, allowing their elimination by surrounding WT cells. Our data are consistent with the notion that NMDAR exerts its effect by regulating metabolic coupling between glycolytic loser and oxidative winner cells, driving loser cells to transfer their carbon fuel (lactate) to their neighbours. Preventing loser cells from ‘donating’ lactate to their neighbours removes fitness disparities and abrogates NMDAR-mediated cell competition.

## Results

### Clonal depletion of NR2 results in the elimination of otherwise viable cells

Genetic studies in *Drosophila* have revealed that tumour-driven cell competition eliminates nearby wildtype neighbours through TNF-dependent cell death, allowing the expansion of oncogene-induced supercompetitor clones^1, 3, 10^. However, the initial mechanism by which cells acquire supercompetitor status to drive cell competition has remained unknown. To identify components that sense fitness disparities during cell competition, we conducted a candidate-based approach in *Drosophila* using hetero-(GFP-marked loser clones among winners – genetic mosaic background) and homotypic (GFP-marked loser clones among losers) competition assays in the developing imaginal wing disc (Figure 1A). Comparison of clonal survival in hetero-versus homotypic genetic backgrounds allows the exclusion of genes that compromise cell viability in general. In *Drosophila*, oncogenic polarity-deficient mutant cells for Discs Large 1 (Dlg1) are eliminated by wild-type neighbours through cell competition^11^. Dlg1 is the highly conserved homologue of mammalian PSD-95 and SAP97, which directly bind to the carboxy-terminal tail of NR2B^12, 13^, a subunit of the *N*-methyl-D-aspartate receptor (NMDAR). NMDAR is best known for its involvement in neuronal activities, including the vast majority of excitatory neurotransmission in the brain, neuronal migration, synaptogenesis, neuronal plasticity, survival, excitotoxicity and lactate-mediated metabolic coupling between neurons and astrocytes^14, 15^. Through these activities NMDA receptors (NMDARs) play roles in long-term memory consolidation, the development of drug addiction, pain perception, and the pathogenesis of neurological disorders^15^. While NMDARs are mainly present in neurons, recent studies indicate that NMDAR subunits are also expressed in non-neuronal tissues^16, 17^. Moreover, high expression of NMDAR is present in various types of cancers, such as human neuroblastoma, breast cancer, small-cell lung cancer and ovarian cancer^18, 19, 20, 21^. Although NMDARs have been extensively studied in the brain, their role in nonneuronal tissues or cancer cells is poorly defined. While there are seven different NMDAR subunits in mammals, *Drosophila* encodes only two NMDAR subunits (NR1 and NR2), which simplifies their study. Our analyses using anti-NR2 antibody staining (Figure S1A) and translational reporter of NR2 (*NR2::GFP*) (Figure S1B), confirmed that NR2 is not only expressed in the *Drosophila* central nervous system^22, 23^ but also in a wide range of developing tissues such as imaginal eye-, and wing discs as well as salivary gland and fat body (Figure S1A–S1C), which is consistent with previously published RNAseq data^17^.

**Figure 1.**
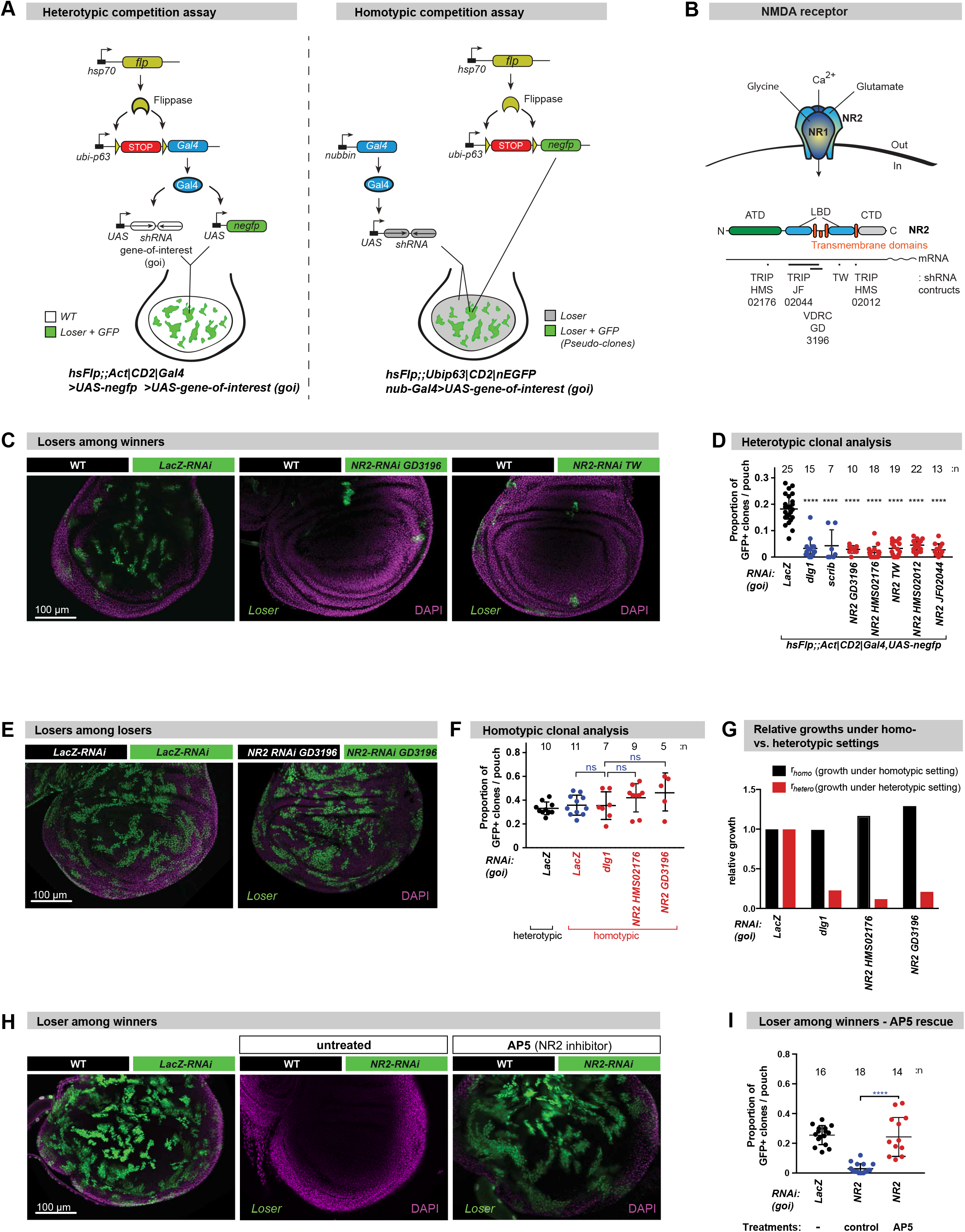
Clonal depletion of *NR2* results in the elimination of otherwise viable cells. **(A)** Schematic representation of the genetic systems used to study cell competition in the *Drosophila* wing pouch. Left panel: Heterotypic genetic system with GFP marking loser clones. Right panel: homotypic genetic system with GFP marking pseudo-clones. **(B)** The *N*-methyl-D-aspartate receptor (NMDAR) is a hetero-tetrameric receptor consisting of two NR1 subunits and two NR2 subunits. Linear representation of the modular amino-terminal domain (ATD, green), ligand-binding domain (LBD, blue), transmembrane domains (yellow) and C-terminal domain (grey). Indicated are the names and position of the respective *NR2*-*shRNAi* constructs used in this study. **(C)** Heterotypic clonal analysis. The indicated genes-of-interest (goi) were knocked down in GFP-marked clones as depicted in a (left panel, see Methods for details). Loser clones are marked by GFP (green). Specific genotypes of the discs shown in these panels, and all subsequent panels, can be found in Extended Data Table. Scale bar 100 μm. **(D)** Quantification of the heterotypic competition assay. Diagram shows the average occupancy of the indicated *RNAi* clones per wing pouch. **(E)** Homotypic clonal analysis. The indicated genes-of-interest were knocked down throughout the wing pouch as described in a (right panel). GFP-marked clones represent pseudoclones. Scale bar 100 μm. **(F)** Quantification of the homotypic competition assay. Diagram shows the average occupancy of pseudoclones in the indicated wing pouches where the respective gene-of-interest were knocked down. **(G)** Relative rates of growth under hetero versus homotypic clonal settings. **(H)** Heterotypic clonal analysis in the presence or absence AP5, an inhibitor of NR2. Scale bar 100 μm. **(I)** Quantification of the AP5 treatment assay. Error bars represent average occupancy of the indicated *RNAi* clones per wing pouch ± SD. \*\*\*\**P*<0.0001, \*\*\**P*<0.001, \*\**P*<0.01, \**P*<0.1 by Mann-Whitney nonparametric *U-*test. n depicts the number of wing discs. See Supplemental Information Table for genotypes.

To study the role of NR2 in cell competition in wing discs, we generated mosaic tissues of two clonal populations. This confronts wild-type cells (WT) with clones of cells in which the gene-of-interest (*goi*) is depleted by RNAi (Figure 1A, left panel). Such RNAi clones were marked with green fluorescent protein (GFP). Interestingly, clonal knockdown of *NR2* (subsequently referred to as *NR2* clones) using five different RNAi constructs (Figures 1B and 1D) resulted in the elimination of the *NR2* clones (Figures 1C and 1D). Likewise, and as previously demonstrated^24, 25^, clonal knockdown of *Dlg1* or *Scribble (Scrib)* (losers among winners) resulted in the elimination of ‘loser’ knockdown-clones (Figure 1D and Figure S2A). In contrast, clonal depletion of *LacZ*, which served as RNAi control, had no effect (Figures 1C and 1D), Importantly, RNAi-mediated depletion of *NR2* or *Dlg1* had no effect on clonal survival under homotypic condition (losers among losers), such as upon tissue-wide *NR2* depletion using *nubbin-Gal4* (Figures 1E and 1F, Figure S2B) or *hedgehog (hh)-Gal4* (Figure S2C) that drive expression of the RNAi constructs in the entire wing pouch or posterior compartment of the *Drosophila* wing imaginal disc, respectively. The observation that *NR2*-depleted cells are lost from the tissue when surrounded by WT cells, but are fully viable under homotypic conditions, demonstrates that clonal reduction of NR2 triggers competitive interactions, resulting in the elimination of otherwise viable cells.

Homotypic clonal analysis demonstrated that the growth rate of *NR2* clones is equivalent to the one of control *LacZ* cells (Figure 1G), highlighting that *NR2* depletion does not impair cell viability or growth in general. Further, treatment with AP5 ((2*R*)-amino-5-phosphonopentanoate), a selective inhibitor of NR2^26^, suppressed the elimination of *NR2* clones in a heterotypic genetic background (losers among winners) (Figures 1H and 1I), phenocopying a homotypic setting. This illustrates that the competitive behaviour between *NR2*-losers and WT-winners is due to a relative difference in NR2 activity among competing clones. AP5-medited global inhibition of NR2 thereby seems to eradicate the “*fitness disparity*” among competing clones. Together, out data demonstrate that depletion of NR2 does not inherently compromise cell viability. However, when surrounded by WT cells, *NR2*-depleted cells are recognized and actively eliminated.

### Elimination of *NR2* loser clones via apoptosis is dependent on the TNF>JNK-signalling axis

To study the elimination process of *NR2-*depleted cells, we examined the possible involvement of the Grindelwald>JNK signalling axis^27^. While NR2-depleted cells were readily eliminated, such loser clones survived upon simultaneous clonal depletion of the *Drosophila* TNF-receptor superfamily member *Grindelwald* (*Grnd*) (*Grnd-RNAi*), *hemipterous (hep)* (*hep-RNAi*) or inactivation of *basket (bsk)* (UAS-*bsk-dominant negative (DN)*), demonstrating that *NR2* clones are eliminated in a Grnd- and JNK-dependent manner (Figure 2A). Consistent with an involvement of JNK signalling, we found intense staining of activated JNK [p-JNK] (Figure 2B), induction of the JNK activity reporter *PucLacZ* (Figure S3A) and high expression level of the JNK target gene *MMP1* (JNK target) in and around *NR2* clones (Figure 2C). JNK signalling in *NR2* clones ultimately resulted in caspase-mediated cell death because expression of the caspase inhibitors p35 and DIAP1 suppressed the elimination of *NR2* clones (Figure 2A). Consistently, we observed cells positive for cleaved caspase staining with anti-cleavedDCP1 (anti-cDCP1) (Figure S3B).

**Figure 2.**
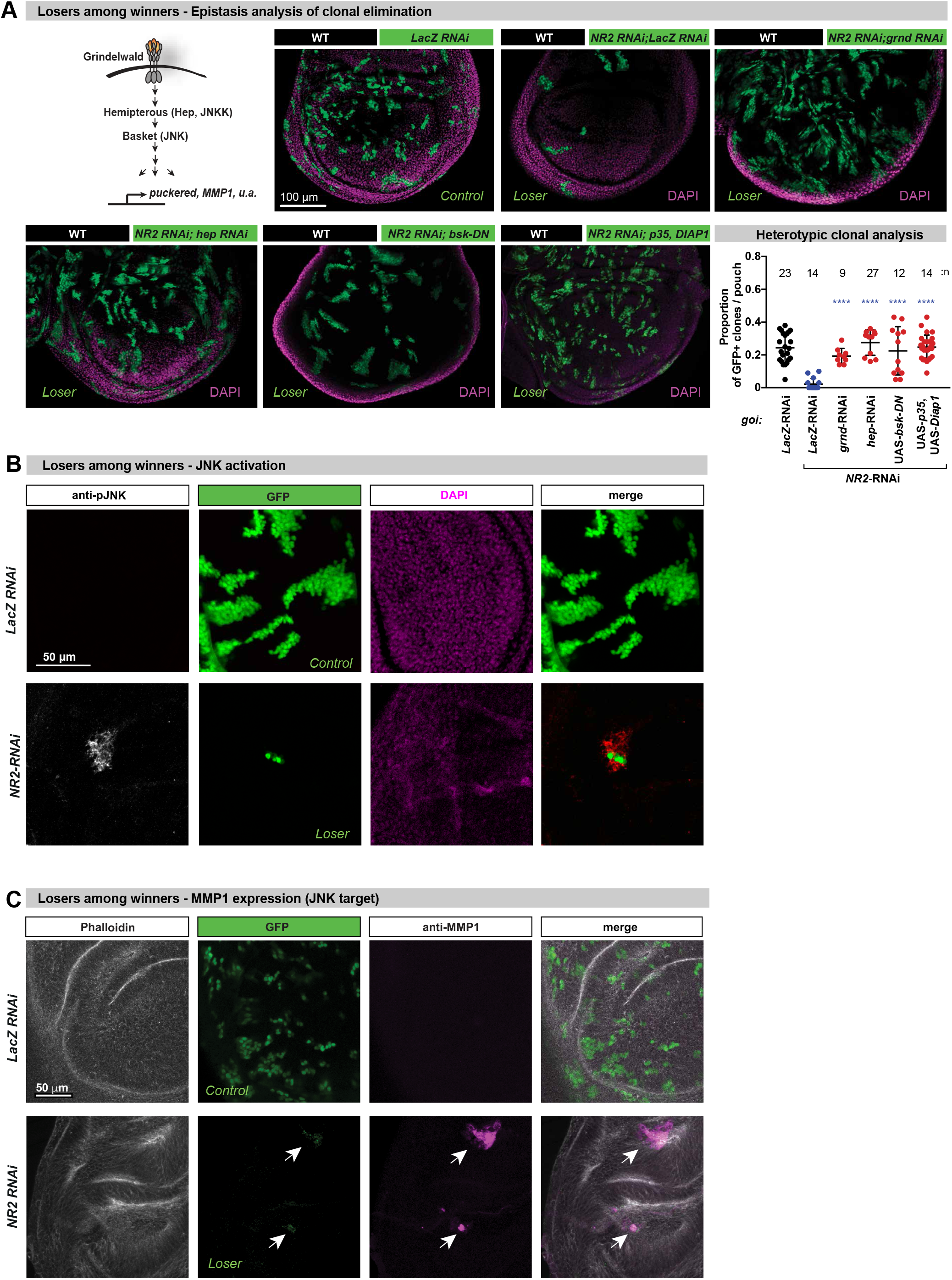
*NR2* loser clones are eliminated by apoptosis via the TNF>JNK signalling axis. **(A)** Heterotypic clonal analysis. Schematic representation of the TNF>JNK signalling pathway. The indicated genes-of-interest (goi) were knocked down or misexpressed (*UAS-bsk-DN*, *UAS-p35,Diap1*) in GFP-marked clones as depicted in Figure 1aA (left panel). Loser clones are marked by GFP (green). Scale bar 100 μm. Quantification of the heterotypic competition assay. Diagram shows the average occupancy of the indicated *RNAi*/overexpression clones per wing pouch ± SD. \*\*\*\**P*<0.0001, \*\*\**P*<0.001, \*\**P*<0.01, \**P*<0.1 by Mann-Whitney nonparametric *U*-test. n depicts the number of wing discs. See Supplemental Information Table for genotypes. **(B)** Confocal images of wing discs that were immunostained with anti-phospho-JNK. Scale bar 50 μm. **(C)** Confocal images of wing discs that were immunostained with Phalloidin and anti-MMP1. Scale bar 50 μm. Supplemental Information Table for genotypes.

### JNK-mediated inactivation of PDH in loser cells induces aerobic glycolysis and lactate production

JNK signalling can lead to activation of the mitochondrial Pyruvate Dehydrogenase Kinase (PDK), which in turn can phosphorylate and inactivate Pyruvate Dehydrogenase (PDH), a key enzyme that catalyses the oxidative decarboxylation of pyruvate to produce acetyl coenzyme A, NADH and CO2^28^. Phospho-dependent inactivation of PDH decreases mitochondrial activity and enhances the conversion of pyruvate to lactate in the cytosol^29^. To test whether depletion of NR2 results in metabolic reprogramming, we monitored the phosphorylation status of PDH. As shown in Figure 3A and Figures S4A–S4C, we detected prominent phospho-PDH (p-PDH) staining in *NR2*-RNAi loser clones. Enhanced p-PDH staining was strictly JNK-dependent as inhibition of JNK signalling abrogated p-PDH staining (Figure 3A). Consistent with PDH inactivation in *NR2*-RNA clones, we noticed a decreased average size of mitochondria (Figure S4D)).

**Figure 3.**
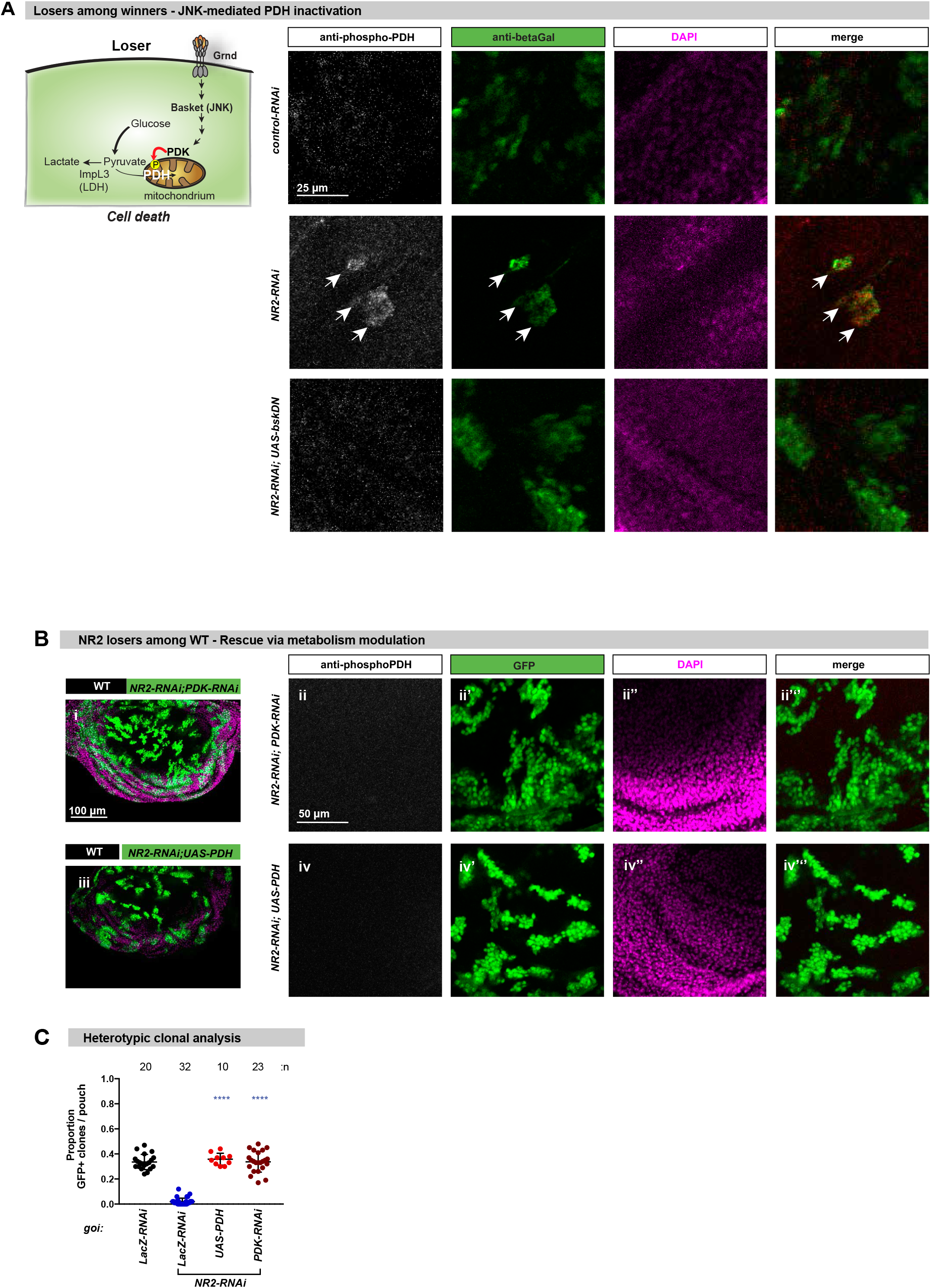
The TNF>JNK>PDK>PDH signalling axis reprograms the metabolism of *NR2* loser clones. **(A)** Schematic representation of the metabolic reprograming of loser cells by the TNF>JNK>PDK>PDH signalling pathway. Phospho-PDH-specific immunostaining of clones expressing genes-of-interest (goi) and UAS-LacZ. LacZ expression is revealed by anti-*β*Gal staining. Scale bar 25 μm. **(B)** Heterotypic clonal analysis. The indicated genes-of-interest (goi) were knocked down or over-expressed (*UAS-PDK-RNAi* or *UAS-PDH*) in GFP-marked clones, and stained for anti-phospho-PDH (ii,iv). Loser clones are marked by GFP (green). Scale bar 100 μm (i,iii) or 50 μm (ii,iv). **(C)** Quantification of the heterotypic competition assay. Diagram shows the average occupancy of the indicated *RNAi*/overexpression clones per wing pouch ± SD. \*\*\*\**P*<0.0001, \*\*\**P*<0.001, \*\**P*<0.01, \**P*<0.1 by Mann-Whitney nonparametric *U-*test. n depicts the number of wing discs. See Supplemental Information Table for genotypes.

Next, we tested the importance of PDK and p-PDH for the elimination of *NR2*-RNAi loser clones. Intriguingly, co-depletion of PDK completely rescued the elimination of *NR2*-loser clones (Figures 3B and 3C) and abrogated the appearance of cleaved DCP1 positive cells (Figure S4E). Likewise, Gal4-driven expression of PDH in *NR2*-loser clones blocked their elimination (Figures 3B and 3C). In both settings, surviving *NR2*-clones were negative for anti-p-PDH staining (Figure 3B). Together, these data indicate that PDK-mediated phosphorylation and inactivation of PHD is required for the death of NR2-loser clones.

### Loser cells die by ‘donating’ their lactate to winners

Since inactivation of PDH results in aerobic glycolysis, we assessed whether metabolic reprogramming of loser cells might contribute to their elimination. During aerobic glycolysis, most glucose carbon is converted to pyruvate via glycolysis and reduced to lactate via lactate dehydrogenase (LDH)^30^, which is then secreted. While lactate exits cells via monocarboxylate transporter (e.g. MCT1) to avoid acidification, it can be recaptured and used as source of energy by neighbouring cells, leading to metabolic compartmentalisation between adjacent cells^4, 31^. To test the involvement of lactate exchange from losers to winners during cell competition, we exposed flies to food containing lactate (L-lactic acid, LLA). We hypothesised that lactate feeding might flatten a putative lactate gradient between losers and winners and hence block cell competition. Intriguingly, lactate feeding blocked the elimination of *NR2*-loser clones (Figures 4A and 4B). Consistent with the notion that lactate transport from *NR2*-losers to winners is important for loser/winner relationships and cell competition, we found that blocking loser cells to produce and transport lactate to winner cells rescued loser cell elimination. Accordingly, concomitant down-regulation of the lactate dehydrogenase *ImpL3* in *NR2* loser clones, like co-knockdown of the lactate transporter *monocarboxylate transporter 1 (Mct1)*, completely rescued *NR2* clone elimination (Figures 4A and 4B). This effect was specific to MCT1, as silencing the putative monocarboxylate transporters *CG13907* or *CG3409* within loser clones had no effect on *NR2*-loser cell elimination (Figure 4B). Pharmacological inhibition of MCT1 also blocked cell competition and the elimination of *NR2* loser clones (Figures 4A and 4B). Consistent with these results, we observed elevated levels of MCT1 in *NR2-RNAi* cells (Figure S5). Together, these data suggest that preventing loser cells from transferring lactate to their neighbours may remove fitness disparities and inhibit cell competition.

**Figure 4.**
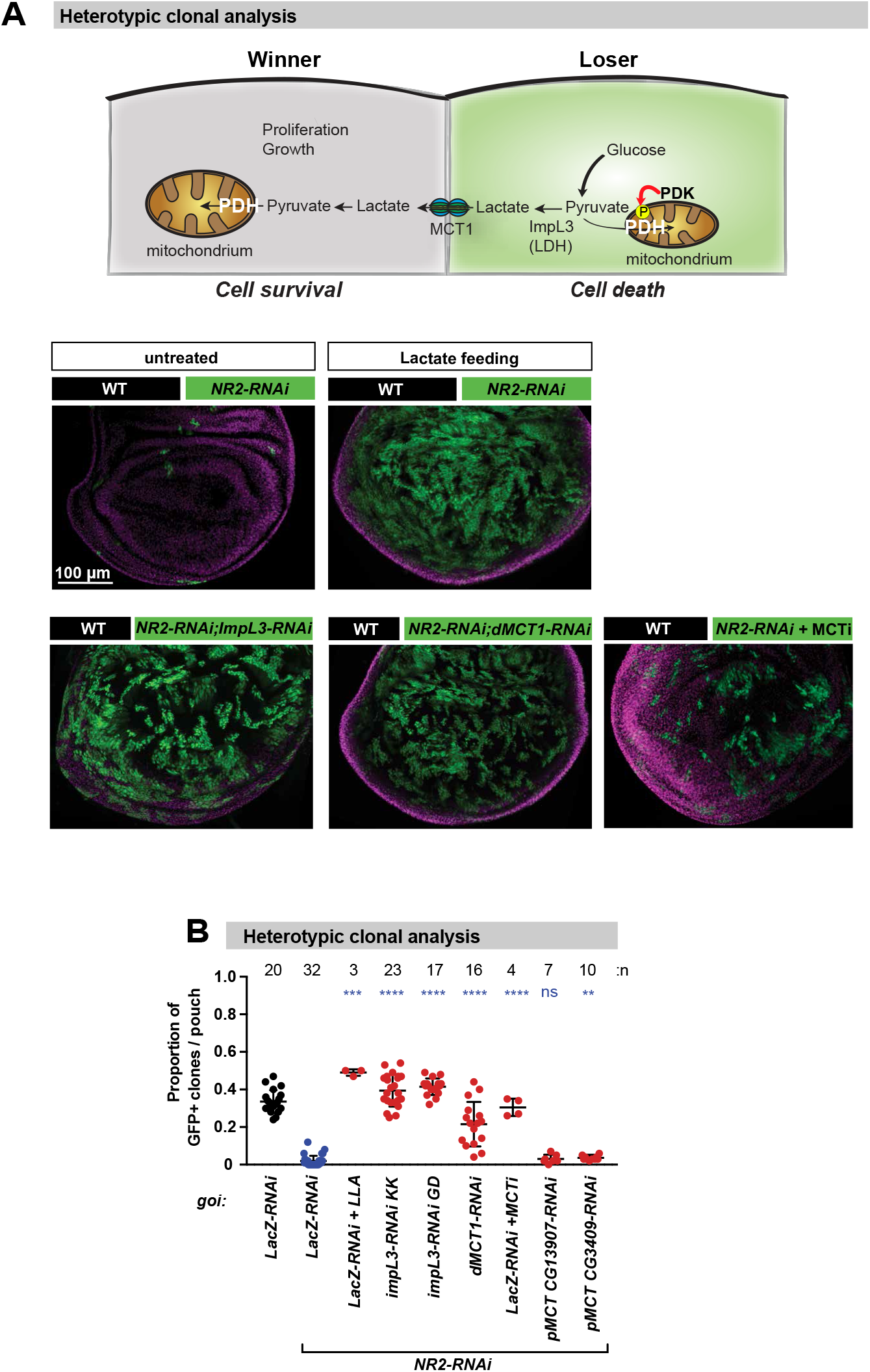
NR2 controls cell competition by regulating lactate-mediated metabolic coupling between winners and losers. **(A)** Schematic summary of our data depicting NR2-assisted regulation of cell competition via lactate-mediated metabolic coupling. Heterotypic clonal analysis. The indicated genes-of-interest (goi) were knocked down and over-expressed in *GFP*-marked clones. For genotypes see Supplementary Data Table. Loser clones are marked by GFP (green). Lactate feeding and treatment with MCT inhibitor was conducted as outlined in the Methods section. **(B)** Quantification of the heterotypic competition assay. Diagram shows the average occupancy of the indicated *RNAi* clones per wing pouch, using the indicated conditions. \*\*\*\**P*<0.0001, \*\*\**P*<0.001, \*\**P*<0.01, \**P*<0.1 by Mann-Whitney nonparametric *U*-test. n depicts the number of wing discs. See Supplemental Information Table for genotypes.

### NR2 is essential for the supercompetitior status of Myc- and RasV12-expressing cells

In mosaic wing imaginal discs, interactions between WT and Myc-or RasV12-expressing cells cause oncogene expressing cells to acquire ‘supercompetitor’ behaviour that increases their fitness and enables them to overtake the tissue by killing their WT neighbours^32, 33^. To test the potential role of NR2 for the suppercompetitor status of Myc-expressing cells, we first examined the levels of NR2 in Myc clones. As shown in Figures 5A and 5B Myc supercompetitor clones exhibited significantly higher levels of NR2 than surrounding wild-type neighbours, both in the wing disc as well as fat body. Moreover, such Myc supercompetitor clones also had higher levels of the ligand for NR2, Glutamate (Figure 5C).

**Figure 5.**
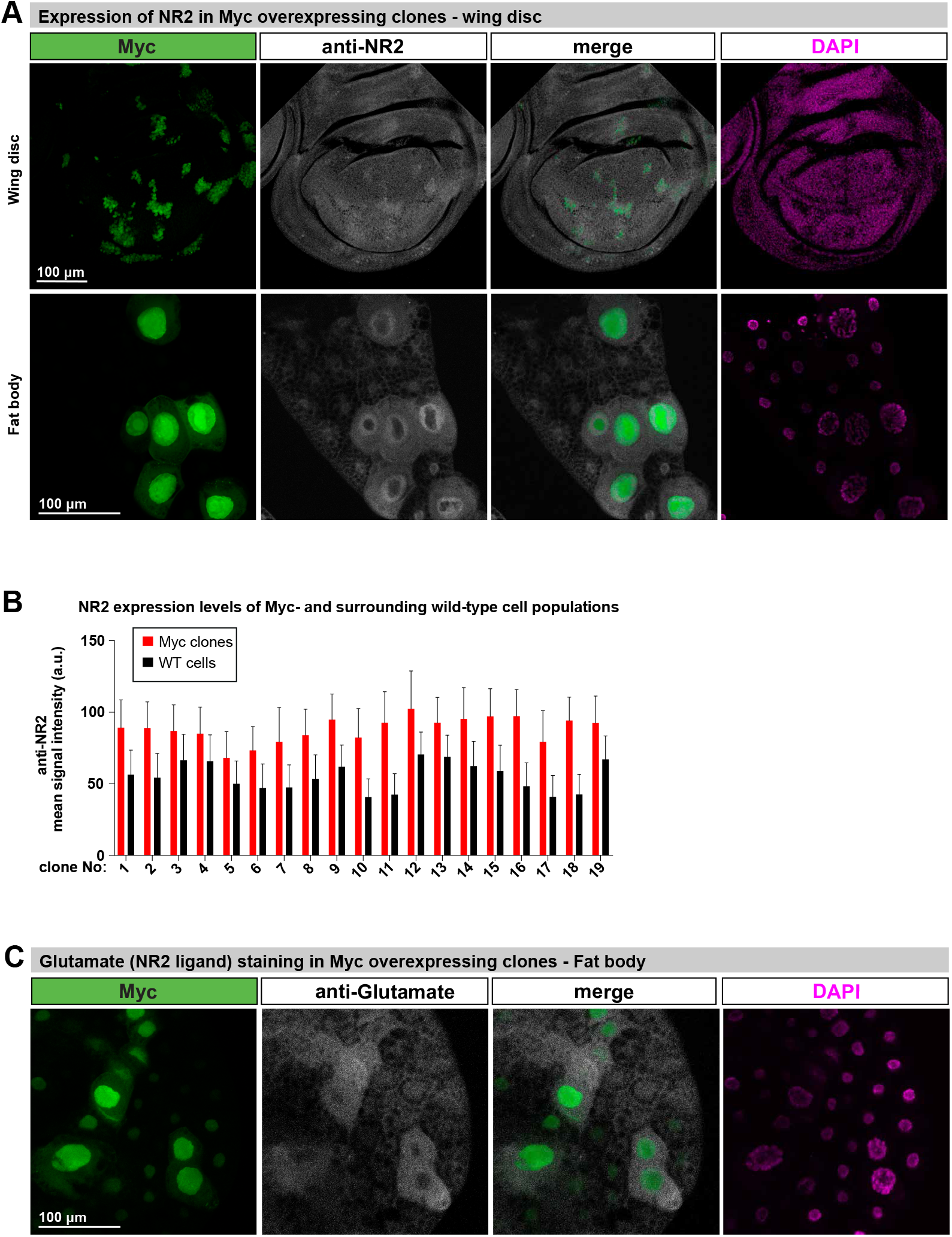
NR2 is upregulated in supercompetitior *Myc*-expressing cells. **(A)** Immunostaining of imaginal wing discs and fat body, GFP-marked Myc overexpressing clones. NR2 expression is revealed by antibody specific to the *Drosophila* NR2 subunit. Scale bars 100 μm. **(B)** Mean signal intensity of anti-NR2 specific staining in Myc expressing cells (red) and immediately adjacent wild type cells (black). **(C)** Anti-Glutamate-specific staining of fat body, Myc overexpressing cells are marked with GFP. Scale bar 100 μm. See Supplemental Information Table for genotypes.

To test whether NR2 contributes to the supercompetitor status of oncogenic Myc, we depleted *NR2* in Myc supercompetitor clones. While clonal expression of oncogenic Myc on its own resulted in large hyperplastic clones, such Myc winners were eliminated when *NR2* was simultaneously knocked down in these clones (Figures 6A and 6C). Likewise, clones expressing oncogenic RasV12 were also eliminated when *NR2* was depleted in RasV12 clones (Figures 6B and 6C). Further, we found that depletion of *NR2* in RasV12 clones not only eliminated RasV12 clones but also rescued pupal lethality, which is associated with the neoplastic growth of RasV12 clones and the development of RasV12 tumours (Figure 6D). Together, these data demonstrate that the supercompetitor status of Myc, and the potential of RasV12 cells to form tumours, critically depends on the presence of NR2. Importantly, Myc- and RasV12-expressing *NR2* clones were eliminated only in heterotypic settings, when surrounded by WT cells. Accordingly, tissue-wide (nub-Gal4) expression of *Myc*;*NR2*-RNAi or *RasV12*;*NR2*-RNAi did not lead to the elimination of GFP+ pseudo-clones, demonstrating that such *Myc*;*NR2*-RNAi and *RasV12*/*NR*2-RNAi clones are intrinsically viable (Figure 6E), but when surrounded by WT cells are eliminated via cell competition and caspase-mediated cell death (Figure 6F).

**Figure 6.**
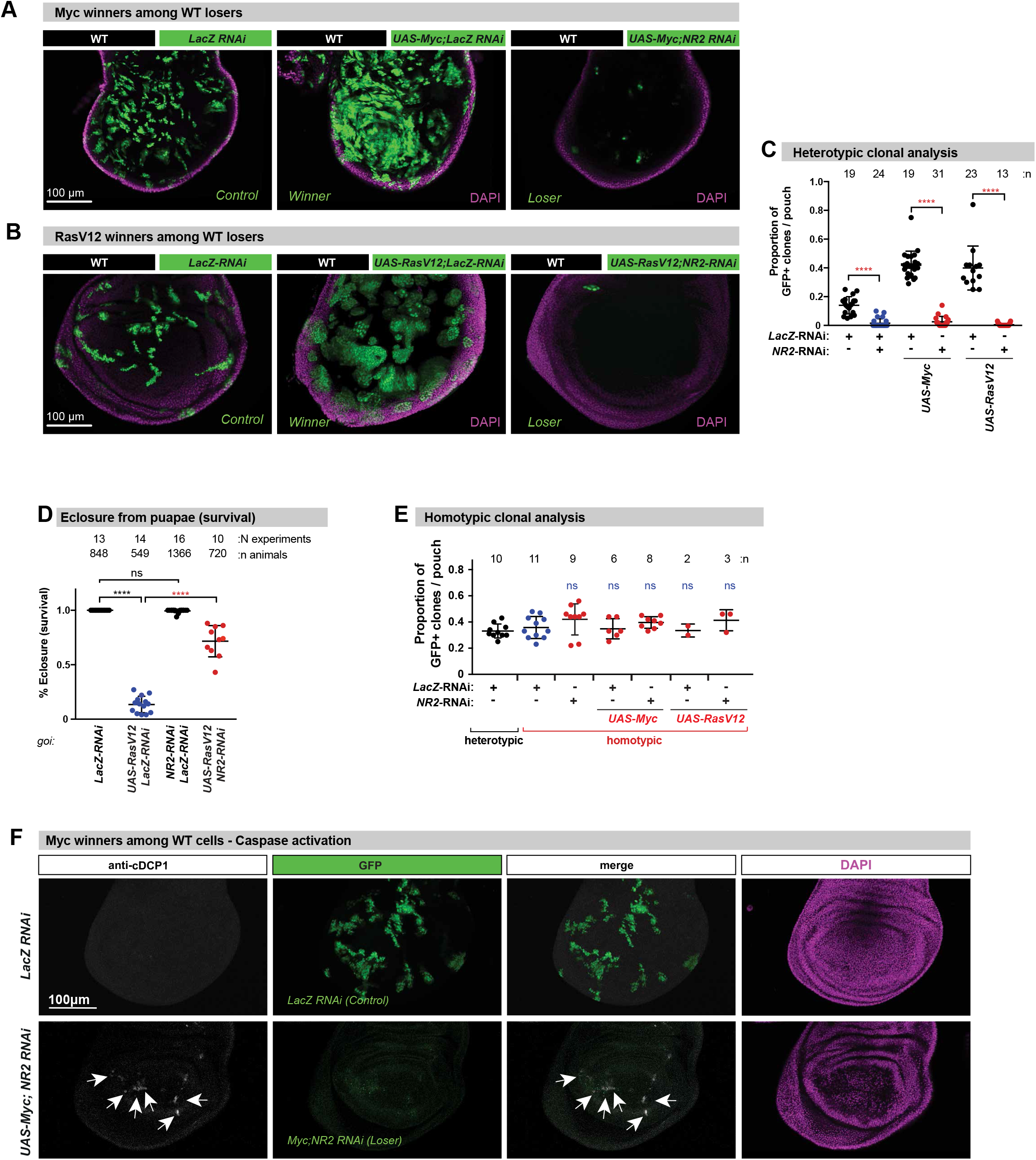
NR2 is essential for the supercompetitior status of *Myc*- or *RasV12*-expressing cells. **(A,B)** Heterotypic supercompetitor clonal analysis. *Myc* (a) *and RasV12* (b) expressing clones are marked by GFP. *LacZ-RNAi* or *NR2-RNAi* were expressed in clones expressing *Myc* or *RasV12* as described in Figure 1A, left panel. **(C)** Quantification of the heterotypic supercompetitor clonal assays. Diagram shows the average occupancy of the indicated clones per wing pouch ± SD. **(D)** Heatshock-mediated induction of *RasV12* clones in the wing pouch caused eclosure-failure and resulted in pupal lethality. Shown are the numbers of eclosed imagoes expressed as the ratio of eclosed imago/total. N: number of independent experiments, n: number of animals. Clones were induced as above. Animals were treated with AP5 as outlined in Methods. **(E)** Homotypic supercompetitor assay. Diagram shows the average occupancy of *Myc* and *RasV12* clones on homotypic background. The indicated genes-of-interest were over-expressed (*Myc* or *RasV12*) and knocked-down (*LacZ* or *NR2*) throughout the wing pouch as described in Figure 1B (right panel). GFP-marked clones represent pseudoclones. Diagram shows the average occupancy of the indicated clones (heterotypic analysis) or pseudoclones (homotypic analysis) per wing pouch ± SD. \*\*\*\**P*<0.0001, \*\*\**P*<0.001, \*\**P*<0.01, \**P*<0.1 by Mann-Whitney nonparametric *U-*test. n depicts the number of wing discs. **(F)** Shown are confocal images of wing discs that were immunostained with anti-cleaved DCP1. Scale bar 100 μm. See Supplemental Information Table for genotypes.

Next, we monitored the p-PDH (metabolic reprogramming) status in WT and Myc or RasV12 supercompetitor cells to assess whether lactate-mediated metabolic coupling might be important for loser/supercompetitor relationships. Under homotypic conditions, no p-PDH staining was apparent in WT cells (Figure S6A and Figure S7 top panel). However, prominent p-PDH staining was detectable in WT cells that were juxtaposed to Myc or RasV12 winner clones (Figures S6B–S6D and Figure S7 middle panel). Interestingly, depletion of *NR2* in RasV12 clones, which renders them losers, lead to strong p-PDH staining in such clones (Figure S7, bottom panel). This suggests that depletion of *NR2* reprograms metabolism via phosphorylation-dependent inactivation of PDH and consequential shut down of pyruvate catalysis, leading to enhanced production and disposal of lactate. According to our model (Figure 4A), this then results in the transfer of lactate from losers to their neighbours, providing them with winner status. Hence, altered metabolism of loser cells may be due to lower NR2 activity. Together, our data suggest that NR2-mediated regulation of metabolism may influences cell competition.

## Discussion

The active elimination of unfit cells via competitive interactions plays an important role for the maintenance of tissue health during development and adulthood^1, 2, 34, 35, 36^. Our data indicate that the activity of NR2 influences the competitive behaviour of epithelia cells and Myc or RasV12 supercompetitors. We find that pharmacological inhibition or genetic depletion of *NR2* reprograms metabolism via TNF-dependent and JNK-mediated activation of PDK, which in turn phosphorylates and inactivates PDH. This causes a shutdown of pyruvate catalysis and results in a switch to aerobic glycolysis. In the absence of functional PDH, pyruvate is reduced to lactate via LDH, and secreted^30^. Disposal of this large carbon pool as lactate serves multiple functions. While lactate exits cells to avoid acidification, it can be recaptured and used as carbon source by other cells, leading to metabolic compartmentalisation between adjacent cells. In normal physiology as well as in murine and human tumours, lactate is an important energy source that fuels mitochondrial metabolism^4, 31^. For example, lactate produced and secreted by astrocytes is transported to neighbouring neurons where it is used as source of energy to support neuronal function^14^. This is akin of the ‘reverse Warburg effect’^6^, also named ‘two-compartment metabolic coupling’ model, where cancer-associated fibroblasts (CAFs) undergo aerobic glycolysis and production of high energy metabolites, especially lactate, which is then transported to adjacent cancer cells to sustain their anabolic need^6^.

Our data suggest that epithelial NMDA receptor activity is responsible for fitness surveillance and to provide clones with oncogenic mutations supercompetitor status. Cells with decreased epithelial NMDA receptor activity become classified as ‘less fit’ and are metabolically reprogrammed to donate their carbon fuel to their neighbours, rendering them ‘fitter’. According to our model, differential NMDAR signalling in adjacent cells triggers lactate-mediated metabolic coupling, and underpins cell competition in epithelia. Consistently, preventing loser cells from ‘donating’ lactate to their neighbours removes the fitness disparity and nullifies cell competition. Likewise, exposure to elevated levels of systemic lactate, such as upon lactate feeding, completely blocks elimination of *NR2* loser clones, most likely because excessive lactate exposure ‘flattens’ the fitness disparity. This suggests that cell competition may be based on NMDAR-mediated metabolic coupling between winners and losers. Importantly, this metabolic coupling only occurs if adjacent cells have differential NMDAR signalling. Consistently, *NR2* losers are only eliminated if they are surrounded by cells with functional NMDAR. This is evident as tissue-wide inhibition of NMDAR by AP5, a selective inhibitor of NR2, blocks elimination of *NR2* loser clones in a heterotypic genetic setting.

Like in flies, NMDAR also seems to be upregulated in genetically engineered mouse models of cancer, where it induces an autocrine glutamate signalling circuit with resultant stimulation of malignancy^21^. Furthermore, the NMDAR pathway is evident in various human tumours and cell lines, and its expression level is associated with poor cancer patient prognosis^21^. We find that NR2 is upregulated in Myc expressing clones and that such cells co-opt epithelial NR2 to promote cell competition, subduing their neighbouring wild-type cells that become re-classified as ‘unfit’. Given that Myc is a major driver of cancer cell growth, and is a hallmark of the disease in nearly seven out of ten cases, blocking Myc’s function would be a powerful approach to treat many types of cancer. However, the properties of the Myc protein itself make it difficult to design a drug against it. We now find that inactivating NR2 in Myc or RasV12 clones kills these cells in a strictly cell competition-dependent manner.

Together, our data suggest that Myc and RasV12 clones critically depend on the activity of NR2 for their supercompetitor status, and that NR2 could serve as an attractive therapeutic target against various types of cancers that have deregulated levels of NR2, with no apparent effects on homotypic wild-type neighbours. This highlights new vulnerabilities of clones with oncogenic mutations, and may pave the way for new anti-cancer treatment approaches that are based on cell competition.

## Acknowledgments

We would like to thank Laura Johnston, Ann-Shyn Chiang, M. Miura for reagents, and Rebecca Wilson, Sidonie Wicky John, Tencho Tenev, Marta Fores Maresma, Celia Domingues, and Katalin Schlett for technical assistance. We would like to thank Bruno Hudry and members of the Meier laboratory for discussions and critical reading of the ms. We would like to apologize to the many authors whose work we could not cite due to space restrictions. A.B. was funded by an EMBO Long Term Fellowship (ALTF-48-2014). Work in the Meier lab is funded by Breast Cancer Now (CTR-QR14-007) and Biological Sciences Research (BBSRC) (BB/L021684/1). We acknowledge NHS funding to the NIHR Biomedical Research Centre.

## Author Contributions

A.B. conceived the study, A.B. and P.M designed the experiments, and A.B. performed the experiments. A.B. and P.M. analysed the data and wrote the manuscript.

## Author Information

The authors declare no competing financial interests. Readers are welcome to comment on the online version of the paper. Correspondence and requests for materials should be addressed to P.M.

## METHODS

### Fly strains

The following strains were used: *UAS-LacZ-RNAi^37^* (from M. Miura), *UAS-dlg1^GD41136^-RNAi* (Vienna *Drosophila* Resource Center, VDRC), *UAS-NR2^TW^-RNAi^22^* (from A.S. Chiang), *UAS-NR2^TRIPJF02044^-RNAi* (Bloomington *Drosophila* Stock Centre, *BDSC*), *UAS-NR2^TRIPHMS02012^-RNAi (BDSC)*, *UAS-NR2^TRIPHMS02176^-RNAi (BDSC)*, *UAS-NR2^GD3196^-RNAi* (*VDRC*), *UAS-grnd^KK109939^-RNAi* (*VDRC*), *UAS-pvf1-RNAi^39038^* (BDSC); *UAS-hep^GD26929^-RNAi* (VDRC), *UAS-bsk-DN* (BDSC 6409), *UAS-p35,UAS-DIAP1 (*PMID: 10675329), pucE69(*puc-LacZ*) (from P. Ribeiro), *UAS-CaMKII^T287D^* (BDSC: 29664), Pdk-RNAi^TRiPGL00009^ (BDSC: 35142), UAS-Pdh (BDSC: 58765), *UAS-Mct1^KK108618^-RNAi* (VDRC), *UAS-impL3^GD31192^-RNAi* (VDRC), *UAS-impL3^KK110190^-RNAi* (VDRC), *UAS-CG13907^KK107339^-RNAi* (VDRC), *UAS-CG3409^GD37139^-RNAi* (VDRC), *UAS-Myc* (BDSC: 9674); *UAS-Ras85D.V12* (BDSC: 64196), *nub-Gal4* (BDSC: 25754), *hh-Gal4* (BDSC:45946 III), *NR2[MI09281-GFSTF.2]* (BDSC: 60566), *UAS-GCaMP3^NLS^* (from C. Schuster) (PMID: 23652205). For the generation of GFP-marked clones the following strains were used: y,w,*hs-flp*;;*Act<CD2<Gal4,UAS-nEGFP^38^* (from T. Neufeld) and *Ubi-p63E<FRT.STOP.FRT<Stringer(nEGFP)* (BDSC: 32251).

### Fly husbandry

Fly stocks were reared on a standard corn meal/agar diet (6.65 % corn meal, 7.15 % dextrose, 5 % yeast, 0.66 % agar, supplemented with 2.2 % nipagin and 3.4 ml/l propionic acid). All experimental flies were kept at 24 °C on a 12 hours light/dark cycle. Flies crosses were set up and kept for 3 days at 24°C. Flies were transferred to fresh vials every day, and fly density was kept to a maximum of 15 flies per vial. For clonal flip-out experiments (homotypic and heterotypic assays), flies were allowed to lay eggs in fresh tubes for 3 hours. 72 hours after egg laying (AEL) larvae were incubated at 37 °C for 10 minutes to induce transgene expression. Following temperature shift, animals were kept at 24 °C. Imaginal wing discs were dissected 48 hours after heat-shock-mediated induction of clones. For the experiment shown in Figure 3D, clones were induced as above. After clone induction, flies were raised for 10 days at 24 °C. The numbers of eclosed imagoes and dead pupae were counted and the ratio of imago/total per vial was calculated.

### Homotypic and heterotypic cell competition assays

Homotypic assay: *hsFlp;nub-Gal4;ubi-p63<FRT.STOP.FRT<Stringer(nEGFP)* flies were crossed to flies carrying the respective *UAS-based transgenes. nub-Gal4* drives expression of the UAS-transgene in the entire wing pouch. Heat shock generates nEGFP-positive ‘pseudoclones’, which have the same genotype as the surrounding wing pouch. Heterotypic assay: *hsFlp;;Act<CD2<Gal4,UAS-nEGFP* flies were crossed to flies carrying the respective *UAS-based transgenes*. Heat shock generates flip-out clones, removing the *<CD2<* cassette, which allows expression of the UAS-based transgenes. Such clones are marked by nEGFP expression.

### Immunohistochemistry

Larval tissues were stained using standard immunohistochemical procedures. Briefly, discs were dissected in PBS, fixed at room temperature for 20 mins in 3.7 % formaldehyde/PBS and washed in 2 % Triton-X100/PBS. All subsequent incubations were performed in 2 % Triton X-100/PBS at 4 °C. Samples were mounted either in Vectashield or Vectashield containing DAPI (Vector Labs). The following primary antibodies were used: mouse anti-dNR2^22^ (1:100, from Ann-Shyn Chiang), mouse anti-NR2B (1:200, Clone 13, Mouse 610416, BD Biosciences), mouse monoclonal anti-N2B (1:200, [S59-36], ARG22230, Arigobio/2BScientific Ltd.), mouse anti-Dlg (1:50, 4F3, DSHB, University of Iowa, Iowa City, IA, USA), mouse anti-GFP (11814460001, Roche), mouse anti-MMP1 (1:50, 5H7B11, DSHB, University of Iowa, Iowa City, IA, USA), rabbit anti-cleaved DCP1 (Asp216) Cell Signaling), rabbit p-JNK (1:100, *Drosophila* p-JNK (Thr183/Tyr185), 81E11, Cell Signaling Technology Inc., Danvers, MA, USA), PHA-555 (Phalloidin-555, A34055, Invitrogen/Molecular probes), mouse polyclonal Anti-MCT1 (ab90582, Abcam), rabbit polyclonal anti-p-PDHE1 (Pyruvate Dehydrogenase E1-alpha subunit (phospho S293), ab92696, Abcam). Note, the phosphorylation site surrounding S296 of human PDHE1 is conserved in *Drosophila* PDHE1, which is encoded by lethal(1)G0334 (CG7010) (e-value 5e-36, query coverage of 99%). Fluorescent secondary antibodies (FITC-, Cy3- and Cy5-conjugated) were obtained from Jackson Immunoresearch.

### Lactate feeding and treatments with inhibitors

Following heat shock-mediated clone induction, larvae were placed on standard food containing L-lactate (30 mM, 71718-10G, Sigma Aldrich), the NR2 antagonist AP5 (5 μM, A8054, Sigma Aldrich) or the MCT1 inhibitor AR-C155858 (100 μM, Bio-Techne (Tocris)) for 48 hrs.

### Quantifications

Imaginal discs were imaged at 20x magnification. Seven Z-stacks were taken for each disc. After imaging, channels were split and maximum Z-projection was analysed. Using DAPI channel images, a line was drawn around the pouch area and measured using ImageJ. The sum areas of GFP-positive clones were measured using the GFP channel. Threshold was adjusted with the Huang autotresholding algorithm to subtract background. The area above the threshold was analysed. Data were collected from at least three independent experiments, and 10 wing discs per genotype and/or condition were analysed, unless stated otherwise. The relative occupancy of GFP positive clones was quantified, and expressed as proportion of GFP-positive clones per pouch (± S.D.).

### Statistics and data presentation

All statistical analyses were carried out using GraphPad Prism7. Comparisons between two genotypes/conditions were analysed with the Mann-Whitney nonparametric two-tailed rank U-test or Pearson’s correlation test. Confocal images belonging to the same experiment, and displayed together, were acquired using the same settings. For the purpose of visualization, the same level and channel adjustments were applied using ImageJ. Of note, all quantitative analyses were carried out on unadjusted raw images or maximum projections. Values are presented as average ± standard deviation (S.D.), *P*-values from Mann-Whitney U-test (non-significant (ns): *P*>0.05; *: 0.05>*P*>0.1; **: 0.1>*P*>0.01; ***: 0.01>*P*>0.001;****: *P*>0.0001 and from Pearson’s analysis α=0.05.

## Supplemental Information

### Figure Legends

**Figure S1.**
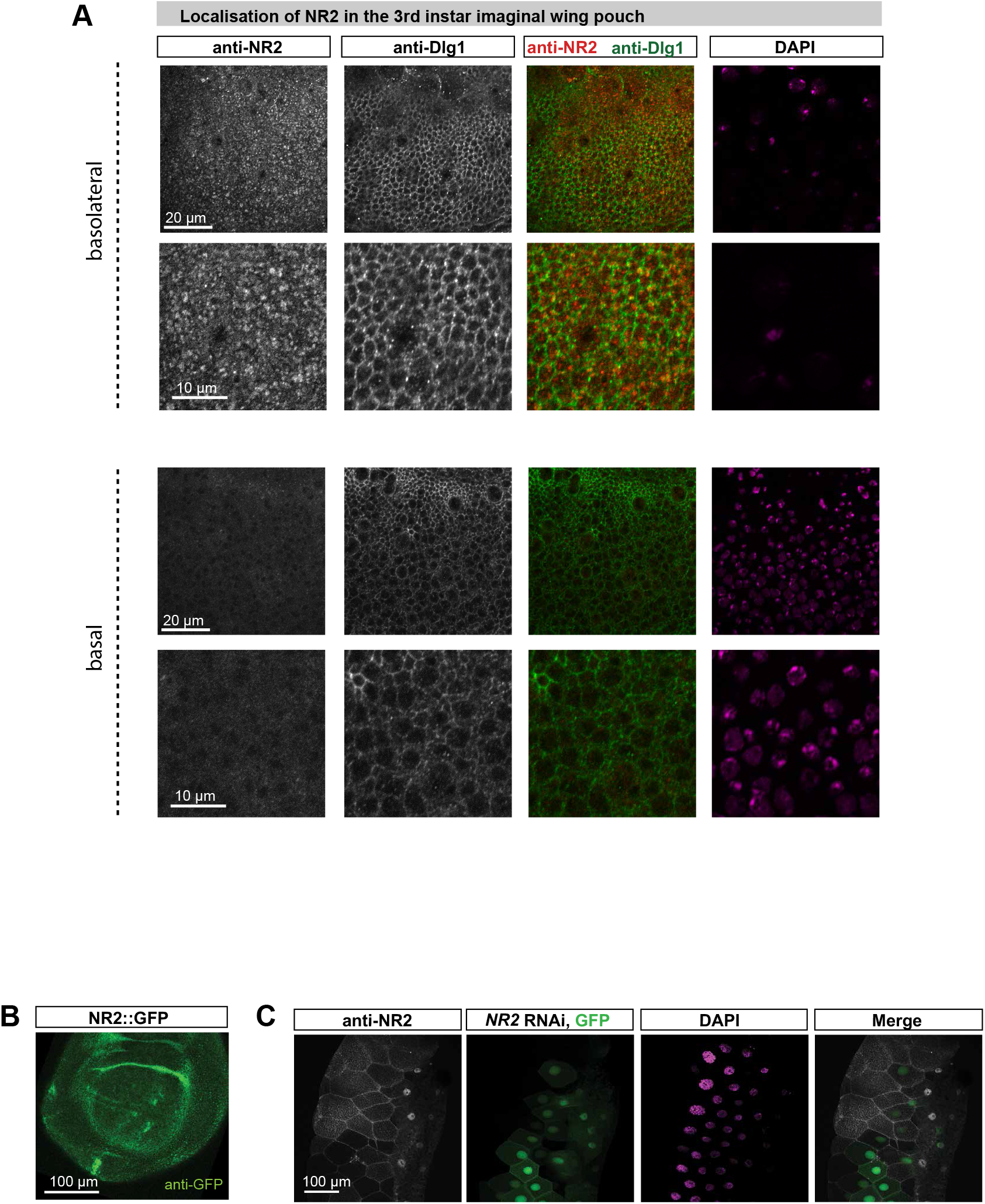
Expression of NR2 in wing discs and salivary glands. **(A)** Localisation and expression of NR2 and Dlg1 in the wing disc of third instar larvae was evaluated using the indicated antibodies. Septate junctions are marked with anti-Dlg1. Scale bars 20 μm and 10 μm. **(B)** Expression analysis of NR2 in the wing disc of third instar larvae. A protein trap line that carries eGFP in frame with NR2 was used to monitor NR2 expression. Shown is a confocal image of a wing disc stained with anti-GFP (NR2). Scale bar 100 μm. **(C)** Expression analysis of NR2 in salivary glands. Antibody staining with anti-NR2 antibodies in salivary glands in which NR2 was downregulated in clones. RNAi clones are marked by GFP. Scale bar 100 μm. See Supplemental Information Table for genotypes.

**Figure S2.**
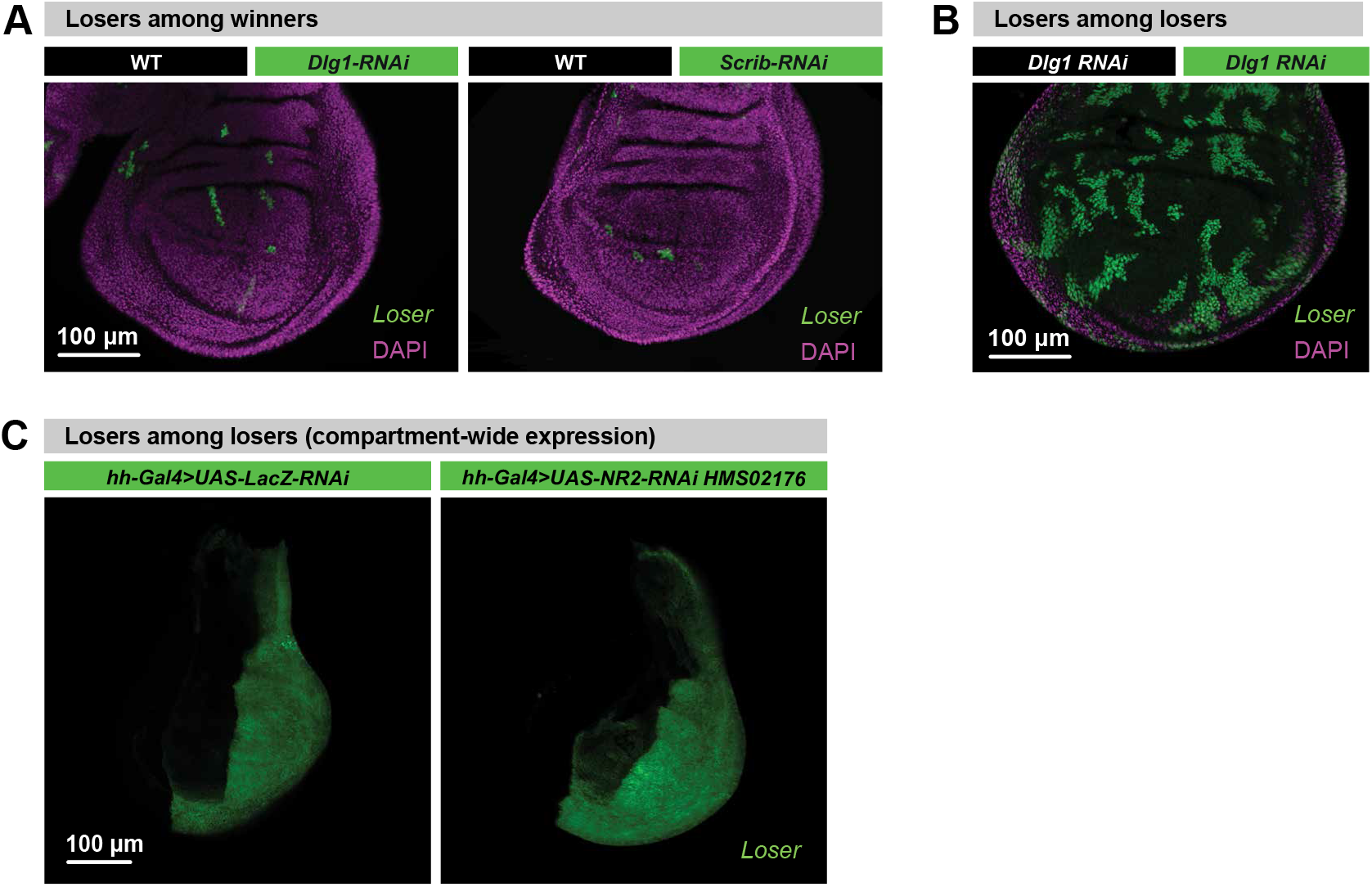
Context-dependent elimination of loser clones. **(A)** Heterotypic clonal analysis. The indicated genes were knocked down in *GFP*-marked clones as depicted in Figure 1D. Scale bar 100 μm. **(B)** Homotypic clonal analysis. *Dlg1* was knocked down throughout the wing pouch as described in Figure 1F. GFP-marked clones represent pseudo-clones. Scale bar 100 μm. **(C)** Homotypic analysis. The indicated genes *were* knocked down throughout the posterior compartment of the wing pouch. Scale bar 100 μm. See Supplemental Information Table for genotypes.

**Figure S3.**
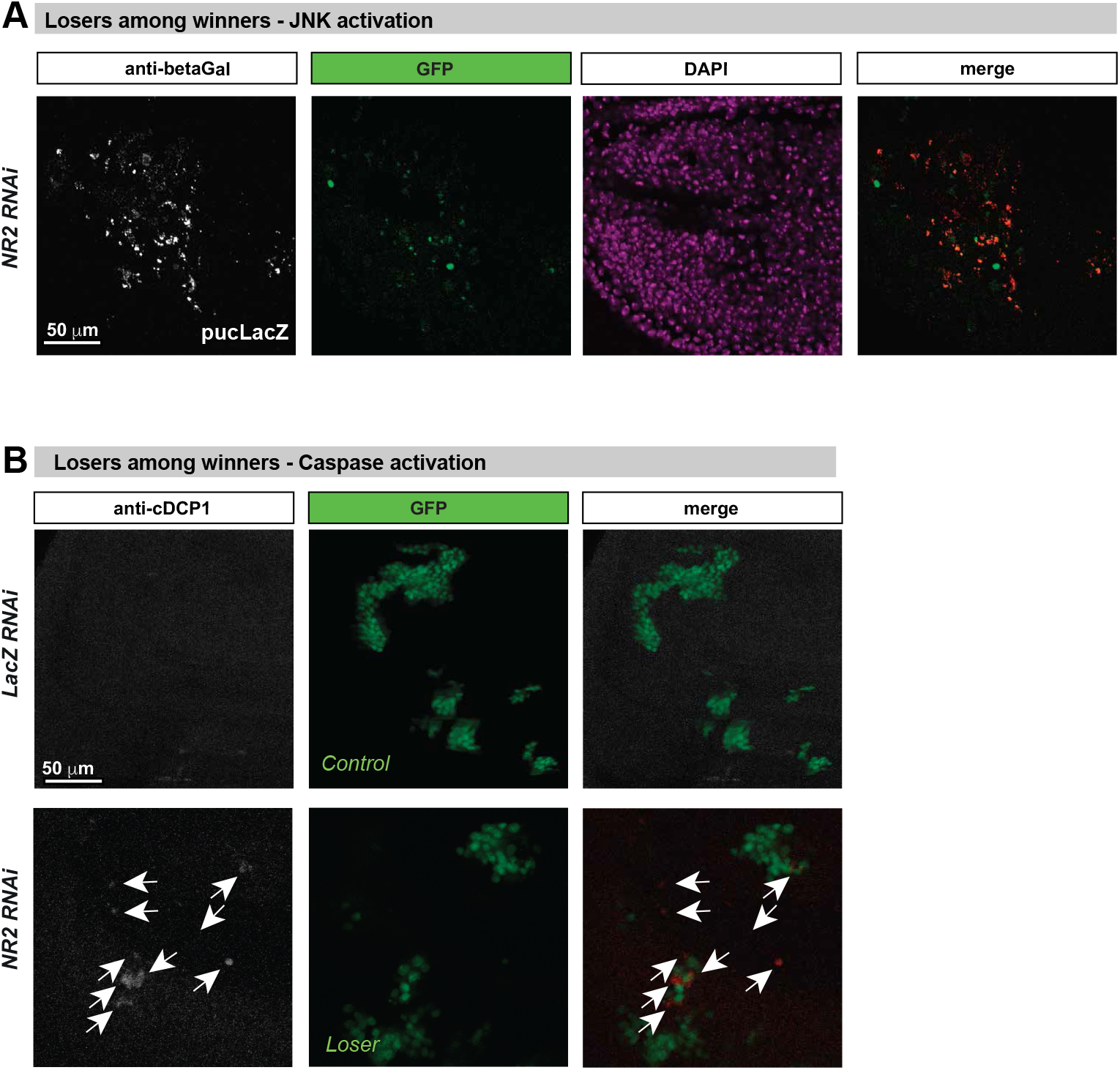
*NR2* loser clones are eliminated by apoptosis via the Eiger>JNK signalling axis. **(A)** Heterotypic clonal analysis of third instar wing discs. *PucLacZ*, expression as a marker of *JNK* activity was revealed with anti-*β*Gal staining. Scale bar 50 μm. **(B)** Confocal images of wing discs that were immunostained with anti-cleaved DCP1. Scale bar 50 μm. See Supplemental Information Table for genotypes.

**Figure S4.**
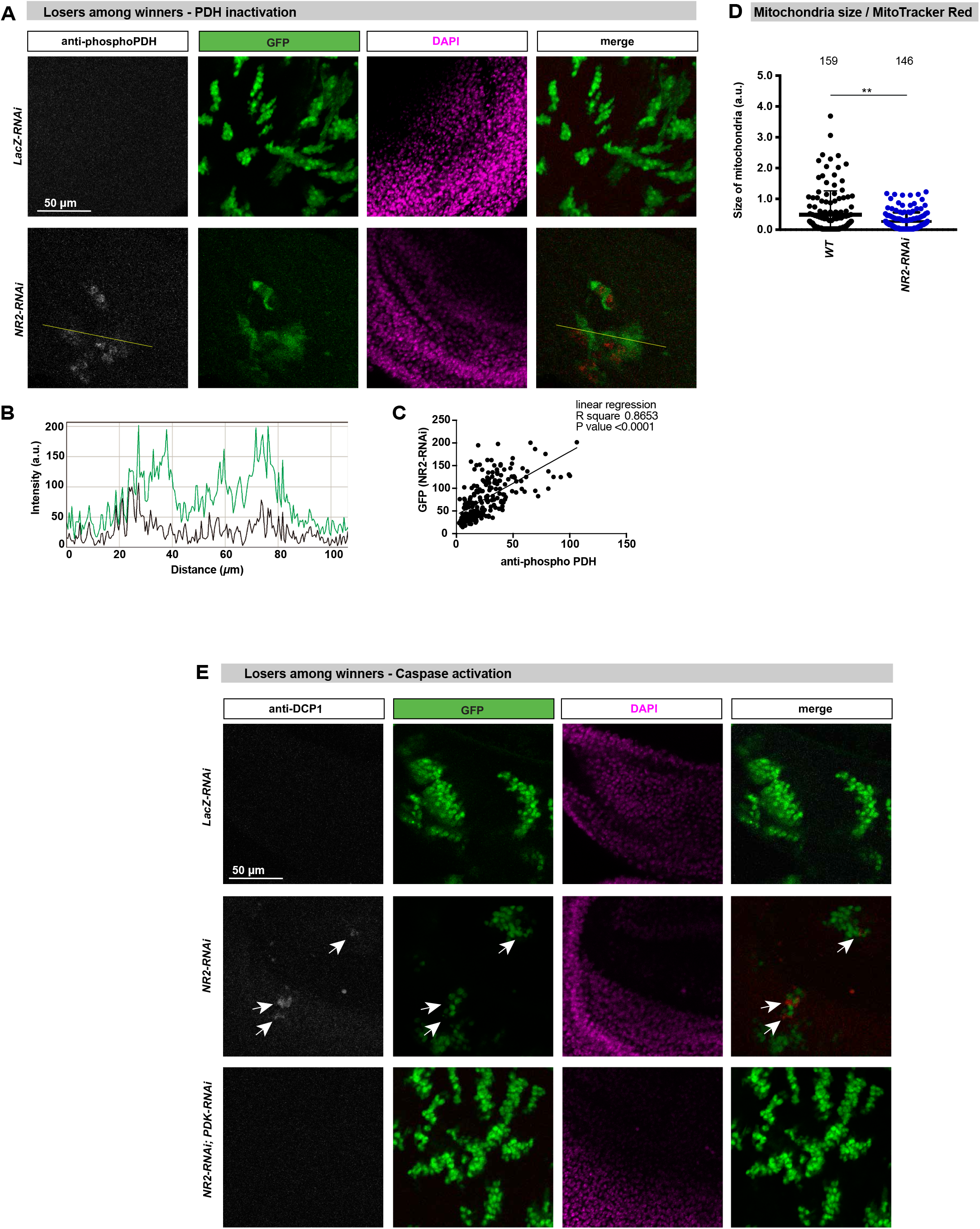
NR2 controls cell competition by regulating lactate-mediated metabolic coupling between winners and losers. **(A)** Confocal images of GFP-marked and anti-phospho-PDH (reading out its inactivation) stained wing pouches. Scale bar 50 μm. **(B)** Fluorescent intensities of phospho-PDH (black) and GFP (green) are measured by ImageJ software at the yellow lines. **(C)** Pearson’s correlation analysis of phospho-PDH and GFP. d, Quantification of the average size of mitochondria stained with MitoTracker Red. **(E)** Confocal images of third instar imaginal wing discs that were immunostained for cleaved DCP1 as a readout of caspase activation and apoptosis. Scale bar 50 μm. See Supplemental Information Table for genotypes.

**Figure S5.**
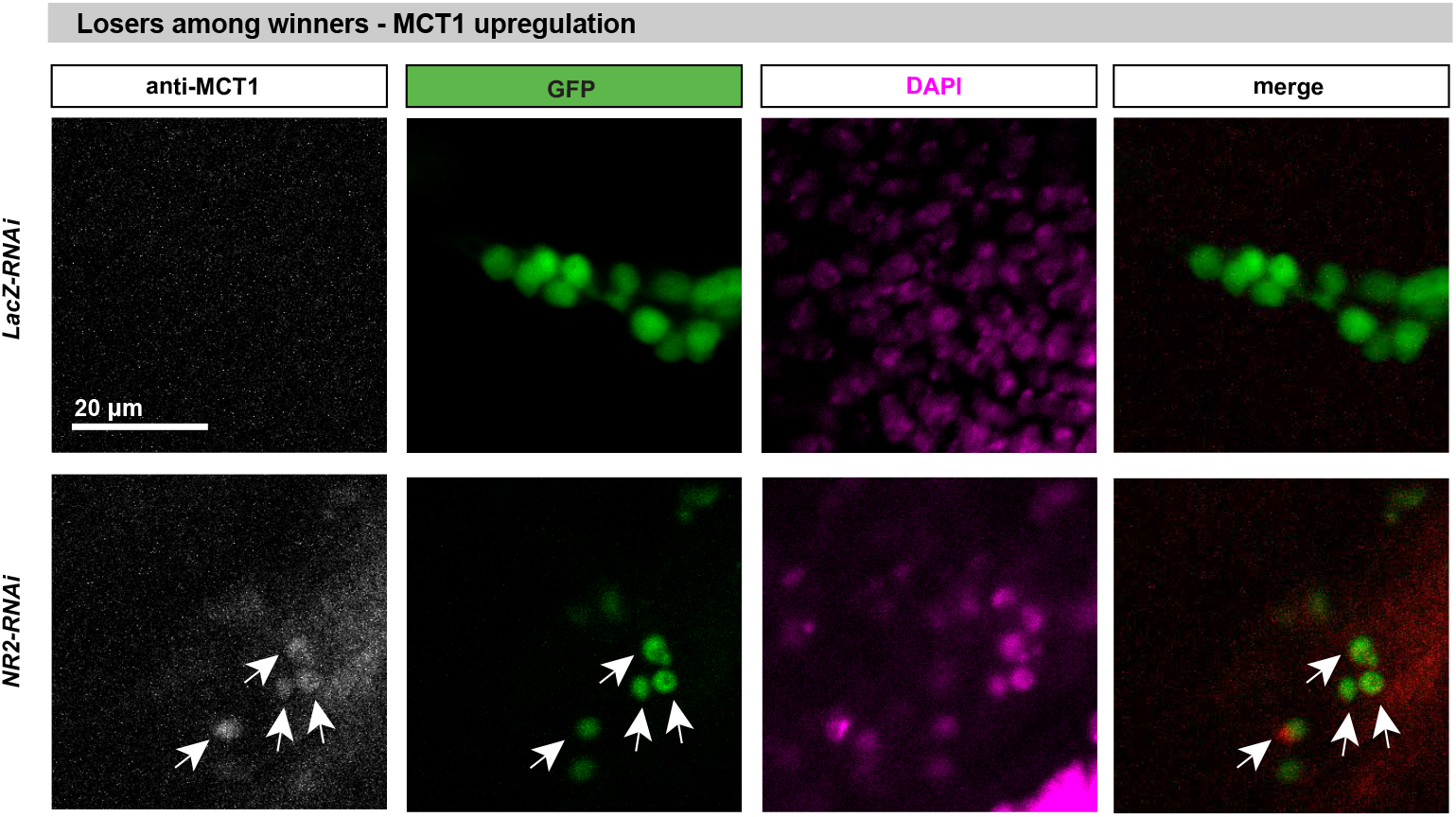
MCT1 is upregulated in *NR2* loser clones. Confocal images of anti-MCT1 stained wing pouches. Scale bar 20 μm. See Supplemental Information Table for genotypes.

**Figure S6.**
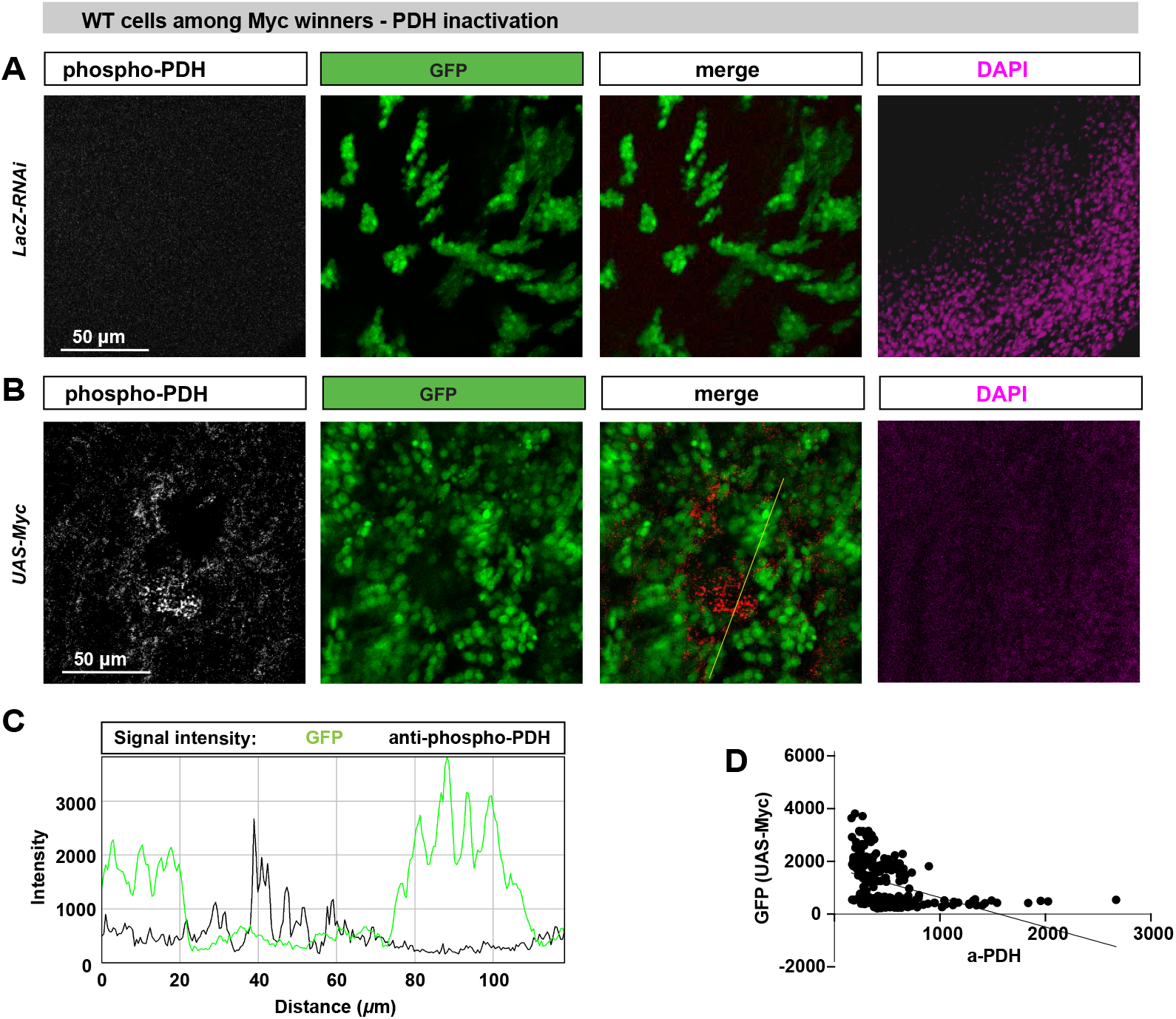
*Myc*-expressing supercompetitior cells reprogram the metabolism of neighbouring wild-type cells. **(A)** Confocal images of dissected wing pouches stained for anti-phospho-PDH (reading out its inactivation). Scale bar 50 μm. **(B)** Fluorescent intensities of phospho-PDH (black) and GFP (green) are measured by ImageJ software at the yellow lines. **(C)** Pearson’s correlation analysis of phospho-PDH and GFP.

**Figure S7.**
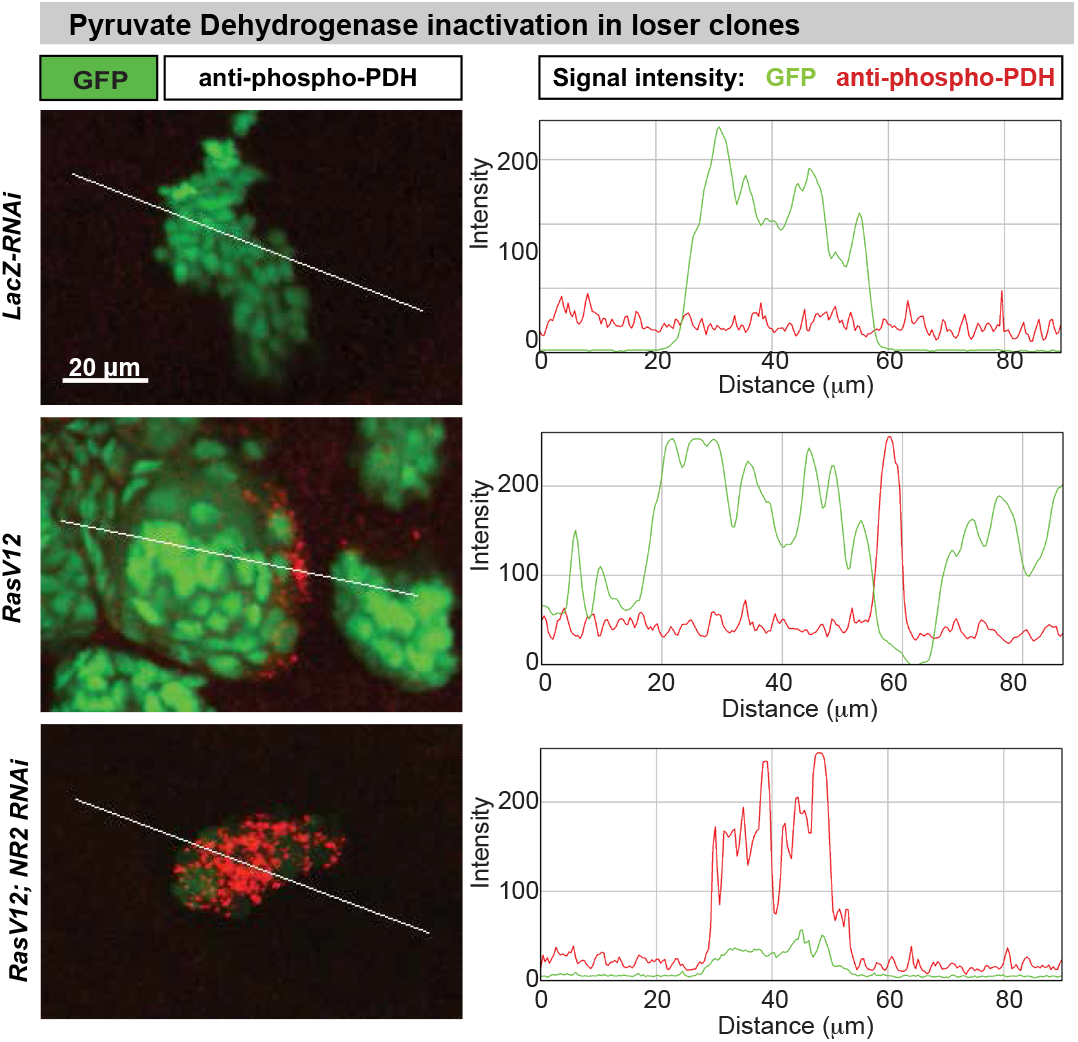
NR2 is essential for the supercompetitior status of *RasV12*-expressing cells. Confocal images of dissected wing pouches stained for anti-phospho-PDH. Scale bar 20 μm. **b**, Fluorescent intensities of phospho-PDH (black) and GFP (green) are measured by ImageJ software at the yellow lines.

## Supplemental Information Table: *Drosophila* genotypes used in this study

### Main Figures

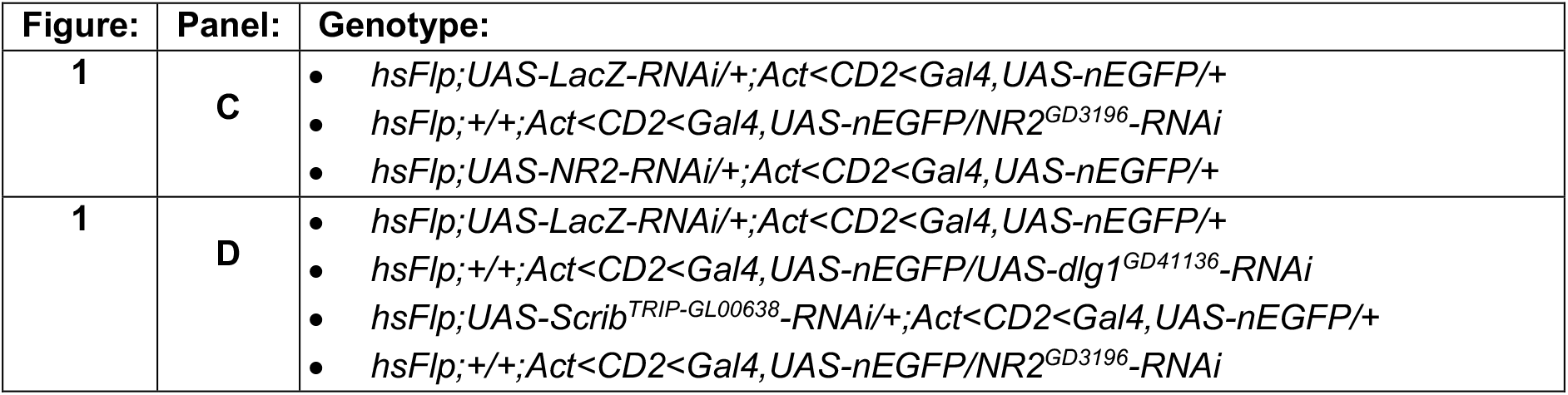
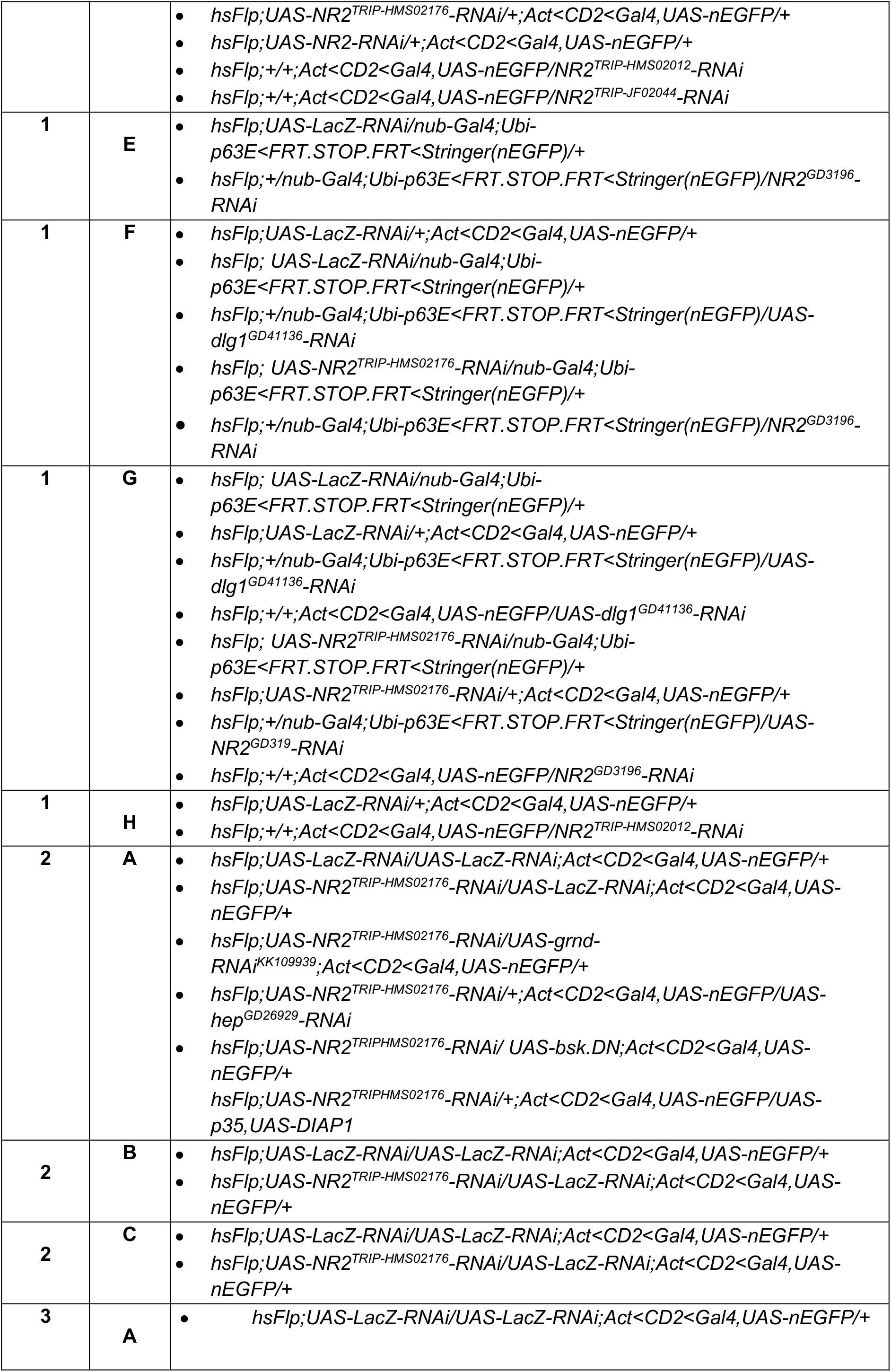
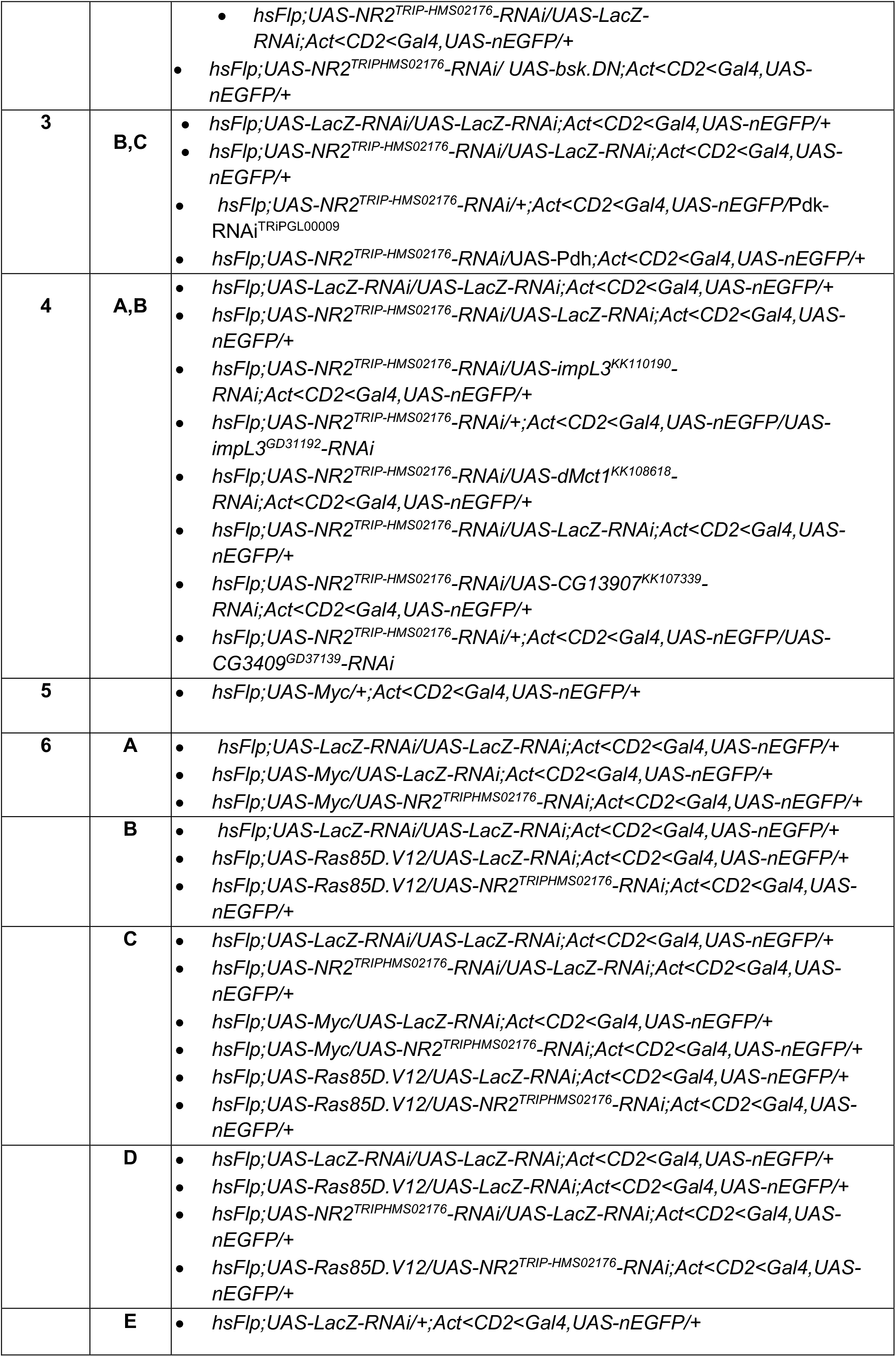
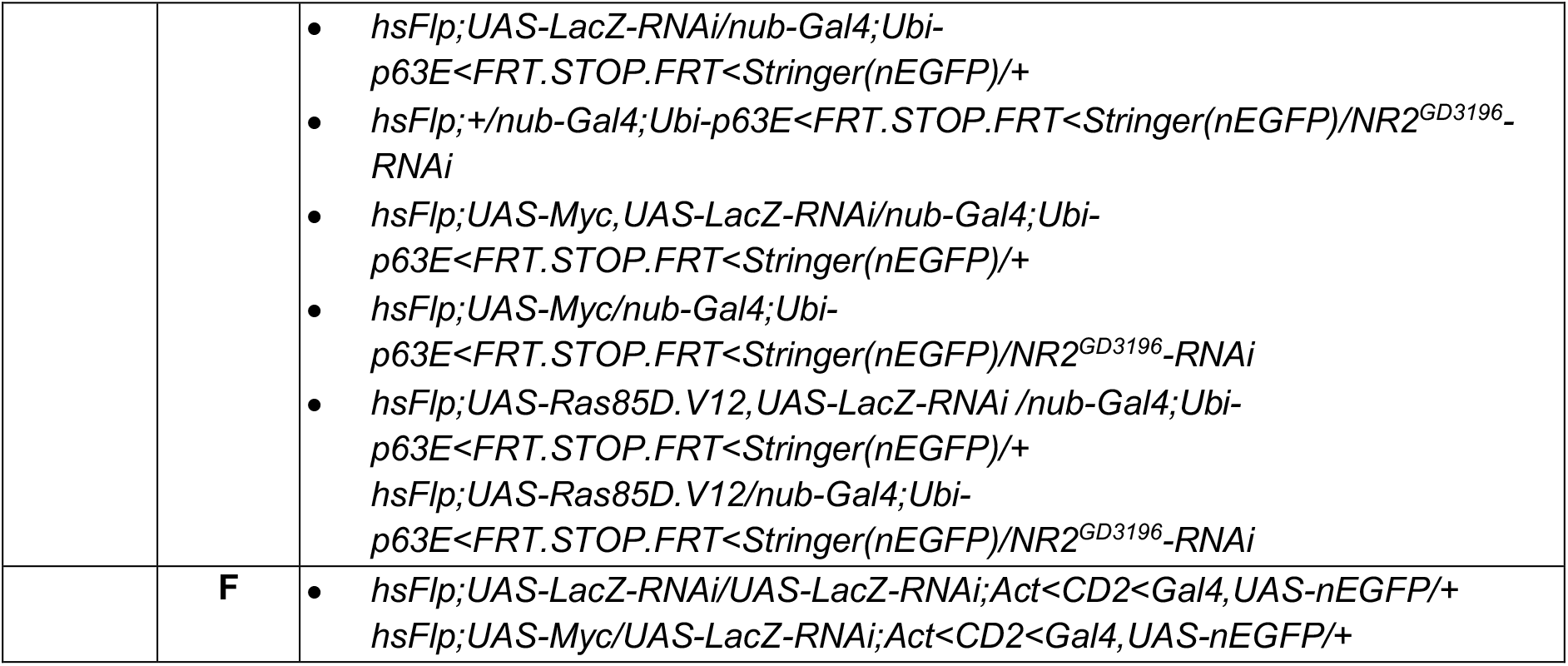

### Supplementary Data Figures

|  |  |  |
| --- | --- | --- |
| **S1** | **A** | • *W1118* |
| **S1** | **B** | • *NR2<sup>MI09281-GFSTF.2</sup>;+;+* |
| **S1** | **C** | • *hsFlp;+/UAS-NR2<sup>TRIP-HMS02176</sup>-RNAi;Act<CD2<Gal4,UAS-nEGFP/+* |
| **S2** | **A** | • *hsFlp;+/+;Act<CD2<Gal4,UAS-nEGFP/UAS-dlg1<sup>GD41136</sup>-RNAi*<br>• *hsFlp;UAS-Scrib<sup>TRIP-GL00638</sup>-RNAi/+;Act<CD2<Gal4,UAS-nEGFP/+* |
| **S2** | **B** | • *hsFlp;+/nub-Gal4;Ubi-p63E<FRT.STOP.FRT<Stringer(nEGFP)/UAS-dlg1<sup>GD41136</sup>-RNAi* |
| **S2** | **C** | • *w1118;UAS-GFP/LacZ-RNAi;hh-Gal4/+*<br>• *w1118;UAS-GFP/ UAS-NR2<sup>TRIP-HMS02176</sup>-RNAi;hh-Gal4/+* |
| **S3** | **A** | • *hsFlp;UAS-NR2<sup>TRIP-HMS02176</sup>-RNAi/UAS-LacZ-RNAi;Act<CD2<Gal4,UAS-nEGFP/pucE69 (puc-LacZ)* |
| **S3** | **B** | • *hsFlp;UAS-LacZ-RNAi/UAS-LacZ-RNAi;Act<CD2<Gal4,UAS-nEGFP/+*<br>• *hsFlp;UAS-NR2<sup>TRIP-HMS02176</sup>-RNAi/UAS-LacZ-RNAi;Act<CD2<Gal4,UAS-nEGFP/+* |
| **S4** | **A** | • *hsFlp;UAS-LacZ-RNAi/UAS-LacZ-RNAi;Act<CD2<Gal4,UAS-nEGFP/+*<br>• *hsFlp;UAS-NR2<sup>TRIP-HMS02176</sup>-RNAi/UAS-LacZ-RNAi;Act<CD2<Gal4,UAS-nEGFP/+* |
| **S5** | **B-D** | • *hsFlp;UAS-NR2<sup>TRIP-HMS02176</sup>-RNAi/UAS-LacZ-RNAi;Act<CD2<Gal4,UAS-nEGFP/+* |
| **S4** | **E** | • *hsFlp;UAS-LacZ-RNAi/UAS-LacZ-RNAi;Act<CD2<Gal4,UAS-nEGFP/+*<br>• *hsFlp;UAS-NR2<sup>TRIP-HMS02176</sup>-RNAi/UAS-LacZ-RNAi;Act<CD2<Gal4,UAS-nEGFP/+*<br>• *hsFlp;UAS-NR2<sup>TRIP-HMS02176</sup>-RNAi/+;Act<CD2<Gal4,UAS-nEGFP/Pdk-RNAi<sup>TRIPGL00009</sup>* |
| **S5** |  | • *hsFlp;UAS-LacZ-RNAi/UAS-LacZ-RNAi;Act<CD2<Gal4,UAS-nEGFP/+*<br>• *hsFlp;UAS-NR2<sup>TRIP-HMS02176</sup>-RNAi/UAS-LacZ-RNAi;Act<CD2<Gal4,UAS-nEGFP/+* |
| **S6** | **A** | *hsFlp;UAS-LacZ-RNAi/UAS-LacZ-RNAi;Act<CD2<Gal4,UAS-nEGFP/+* |
| **S6** | **B-D** | *hsFlp;UAS-Myc/UAS-LacZ-RNAi;Act<CD2<Gal4,UAS-nEGFP/+* |
| **S7** |  | *hsFlp;UAS-LacZ-RNAi/UAS-LacZ-RNAi;Act<CD2<Gal4,UAS-nEGFP/+*<br>*hsFlp;UAS-Ras85D.V12/UAS-LacZ-RNAi;Act<CD2<Gal4,UAS-nEGFP/+*<br>*hsFlp;UAS-Ras85D.V12/UAS-NR2<sup>TRIP-HMS02176</sup>-RNAi;Act<CD2<Gal4,UAS-nEGFP/+* |

